# Prediction of microscopic metastases in patients with metachronous oligo-metastases after curative treatment of Non-Small Cell Lung Cancer

**DOI:** 10.1101/693747

**Authors:** H.B. Wolff, L. Alberts, E.A. Kastelijn, N.E. Verstegen, S.Y. El Sharouni, F.M.N.H. Schramel, R. Vos, V.M.H. Coupe

**Affiliations:** Department of Epidemiology and Biostatistics, Amsterdam Public Health Research Institute, Amsterdam UMC, Vrije Universiteit Amsterdam, De Boelelaan 1118, 1081 HZ, Amsterdam, The Netherlands; Department of Pulmonology, St. Antonius Hospital, Nieuwegein, the Netherlands; Department of Radiation Oncology, Amsterdam UMC, Vrije Universiteit Amsterdam, De Boelelaan 1117, Postbox 7057, 1007 MB, Amsterdam, The Netherlands; Department of Radiation Oncology, University Medical Centre Utrecht, Utrecht, the Netherlands; Department of Methodology & Statistics, Maastricht University Medical Centre, The Netherlands

**Keywords:** Oligo-recurrence, Non-Small Cell Lung Cancer, simulation, tumour growth, prediction model

## Abstract

Metachronous oligo-metastatic disease is variably defined as one to five metastases detected after a disease-free interval and treatment of the primary tumour with curative intent. Oligo-metastases in non-small cell lung cancer (NSCLC) are often treated with curative intent. However additional metastases are often detected later in time, and 5-year survival is low. Burdensome surgical treatment in patients with undetected metastases may be avoided if patients with high versus low-risk of undetected metastases can be separated.

Because there is no clinical data on undetected metastases available, a microsimulation-model of the development and detection of metastases in 100.000 stage I NSCLC patients with a controlled primary tumour was constructed. The model uses data from the literature as well as patient-level data. Calibration was used for unobservable model parameters. Metastases can be detected by a scheduled scan, or an unplanned scan when the patient develops symptoms. The observable information at time of detection is used to identify subgroups of patients with different risk of undetectable metastases. We identified size and number of detected oligo-metastases, as well as presence of symptoms to be the most important risk predictors. Based on these predictors, patients could be divided into a low-risk and a high-risk group having a model-based predicted probability of 8.1% and 89.3% to have undetected metastases, respectively.

Currently, the model is based on a synthesis of literature data and individual patient-level data that was not collected for the purpose of this study. Optimisation and validation of the model is necessary to allow clinical usability. We describe the type of data that needs to be collected to update our model, as well as the design of such validation study.

## Introduction

Metachronous oligo-metastatic disease is variably defined as one to three or one to five metastases detected after a progression-free interval and a controlled primary tumour.(1–4) Oligo-metastases in non-small cell lung cancer (NSCLC) are often treated with curative intent, but additional metastases are commonly detected later in time, and 5-year survival is low in this group of patients.(5) These patients are assumed to have had poly-metastatic disease at time of the oligo-recurrence, with most metastases below the detection threshold of the scan. In this paper we will refer to oligo-metastatic disease with additional undetected metastases as “oligo+”, and without undetected metastases as “oligo-”. Currently, clinicians have no tools available that can distinguish “oligo+” from “oligo-“ patients.(6–8)

Oligo+ patients have no survival benefit from curative therapy. Furthermore, such therapies can result in adverse events or even treatment related mortality.(9, 10) Therefore, it is vital to develop tools to distinguish between oligo+ and oligo-patients, that is, identify predictors that allow for such distinction.

There are two important obstacles for the development of such a tool. Firstly, the true outcome of interest is the presence of unobserved metastases, an outcome that is by definition unknown. Therefore, most studies use surrogate endpoints such as prognostic factors for survival.(11–13) These studies include factors such as patients’ age and performance score into their models, which may select a subgroup of both curable and incurable patients that live relatively longer, or administration of chemotherapy, which can only improve the survival of incurable patients. Those factors are good predictors when the focus is on prognosis in a mixed group of patients with oligo+ and oligo-.

A second obstacle is that there are difficulties with patient accrual in randomized controlled trials.(14) Currently, most studies on oligo-metastases are retrospective studies, with small groups of patients. As a result, these studies are susceptible to selection bias, and levels of evidence are often weak.(15–18)

To tackle both obstacles, we have chosen to construct a microsimulation-model of the development and detection of metastases in stage I NSCLC patients. Microsimulation-models have the advantage that all available evidence can be synthesized to calibrate unobservable parameters, such as the presence of undetectable metastases. In our model, curatively treated patients may develop a number of metastatic lesions that grow exponentially over time. The microsimulation-model keeps track of the growth all metastases within one patient, the number of metastases detected either by surveillance or by symptoms, and recurrence-free survival.

We used the microsimulation-model to construct a simulated patient-level dataset, in which individuals were classified as having detected oligo-metastases or poly-metastases, with the former being subdivided into an oligo− and oligo+ group. Subsequently, we used the simulated dataset to identify clinically observable patient characteristics that can predict presence of undetected metastases.

We describe the development of the microsimulation-model and the underlying evidence. In addition, we present the simulated dataset and the identification of observable patient characteristics that are related to the risk of having undetected recurrences. Finally, we discuss requirements for future validation of our findings and identification of additional predictors, such that identification of patients that benefit from curative treatment of oligo-metastases may be improved.

### Definitions used

**curative treatment:** treatment focused on removing or destroying tumours locally, with the aim to make the patient disease free.

**metastases**: All tumours originating from the primary tumour or another metastasis.

**recurrences**: All metastases detected on a scan.

**oligo-metastasis:** Several definitions have been used in the literature. In this paper oligo-metastasis is defined as 3 or less detected recurrences within one patient, with a controlled primary tumour.

**poly-metastasis**: Several definitions have been used in the literature. In this paper poly-metastasis is defined as 4 or more metastases detected within one patient.

**metachronous oligo-metastasis:** The oligo-metastasis is detected after curative treatment of the primary tumour, contrary to synchronous oligo-metastasis.

**undetectable metastases**: In this paper: all (microscopic) metastases of a size below the detection threshold.

**RFS:** Recurrence-free survival; The proportion of patients that remain recurrence-free after curative treatment of the primary tumour.

**PFS:** Progression-free survival; The proportion of patients that have no additional recurrences detected after curative treatment of oligo-metastases.

**oligo+:** A patient with oligo-metastase(s) with additional undetectable metastases, who would be classified as poly-metastatic if the true number of metastases would be known.

**oligo-:** A patient with oligo-metastase(s) without additional undetectable metastases.

**VDT:** Volume Doubling time; The time required for the tumour to double in volume.

## Methods

### Concept of the microsimulation-model

Curative treatment of oligo-metastases has a pooled 16% 5-year Progression Free Survival (PFS).(6–8, 19, 20) This low PFS is assumed to be caused by undetected metastases that already existed at time of treatment of the primary tumour (Fig 1). The purpose of the micro-simulation model is to test which observable patient characteristics during detection of oligo-metastases may be predictive for the presence of undetected metastases, by synthesizing the available evidence on the growth of tumours, recurrence patterns and methods of detection using current theory and available data.

**Fig 1.**
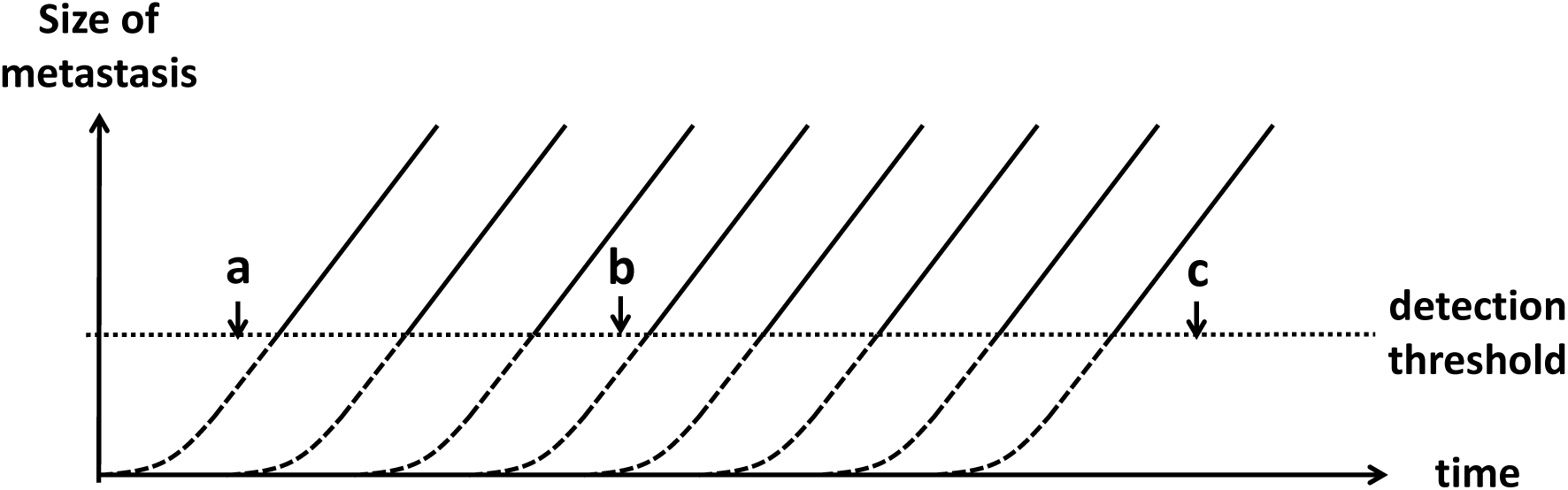
Growth of metastases in time. Metastases are considered undetectable when their size is below the detection threshold (dashed lines), and become detectable above the detection threshold (solid lines). If a patient is scanned at time ‘a’, all metastases are invisible, and this patient would be considered to be ‘recurrence-free’. At time ‘b’, three recurrences would be visible on the scan. This is defined as an oligo-metastasis, even though there are several metastases under the detection threshold (oligo+). At time ‘c’, 8 recurrences are visible on the scan, which is defined as poly-metastatic disease.

The microsimulation-model was developed in C++ and describes the growth and detection of metastases in individual patients. The model stores all patient-specific features and outcomes in comma-separated values files for further analysis. Microsoft Excel Professional plus 2016 was used to create the figures. Code is available upon request.

Certain assumptions and simplifications had to be made, to allow modelling of cancer progression and metastases, without making the model overly complex for the research question that we aim to address (box: model assumptions). The model contains both directly observable parameters as well as unobservable parameters. The observable parameters were directly estimated based on patient data and literature (Table 1). Calibration against observable output variables was used to estimate the unobservable (hidden) parameters, such as the number of undetectable metastases (Table 1).

**Table 1.**
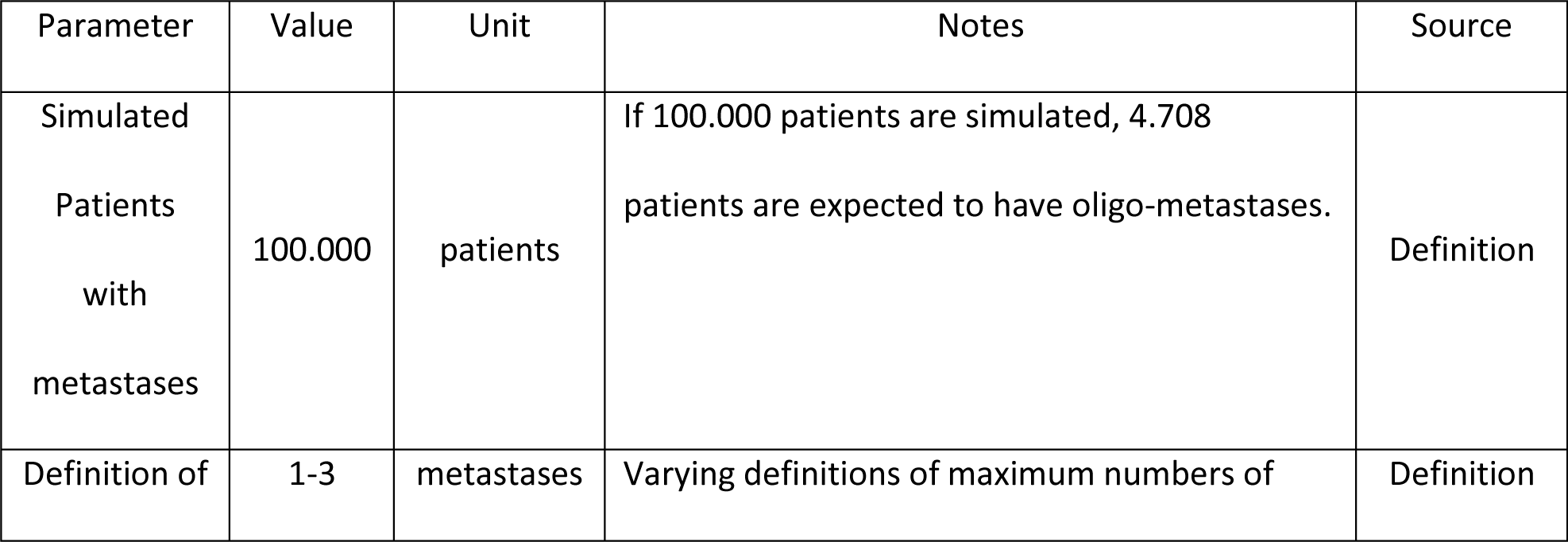
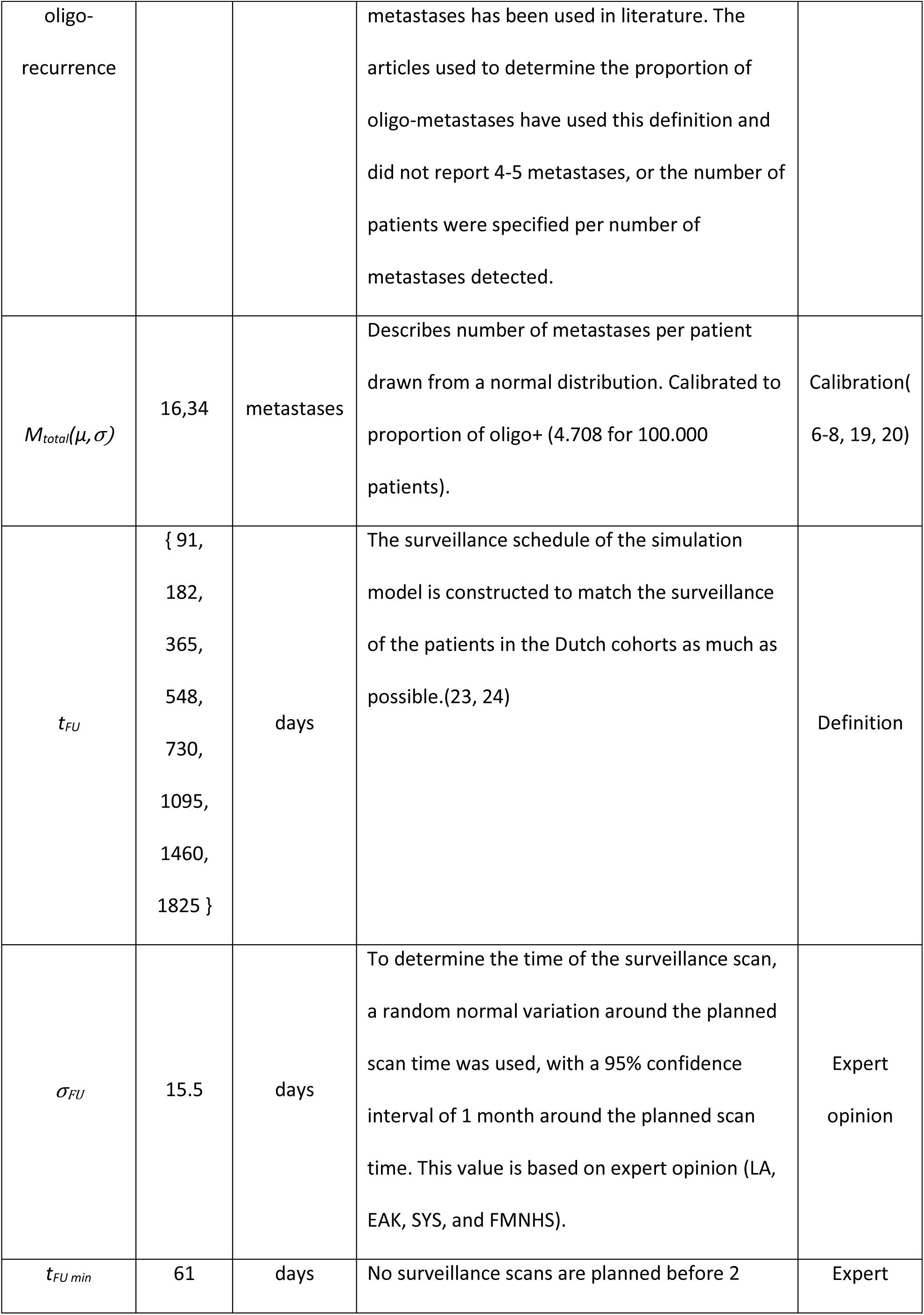
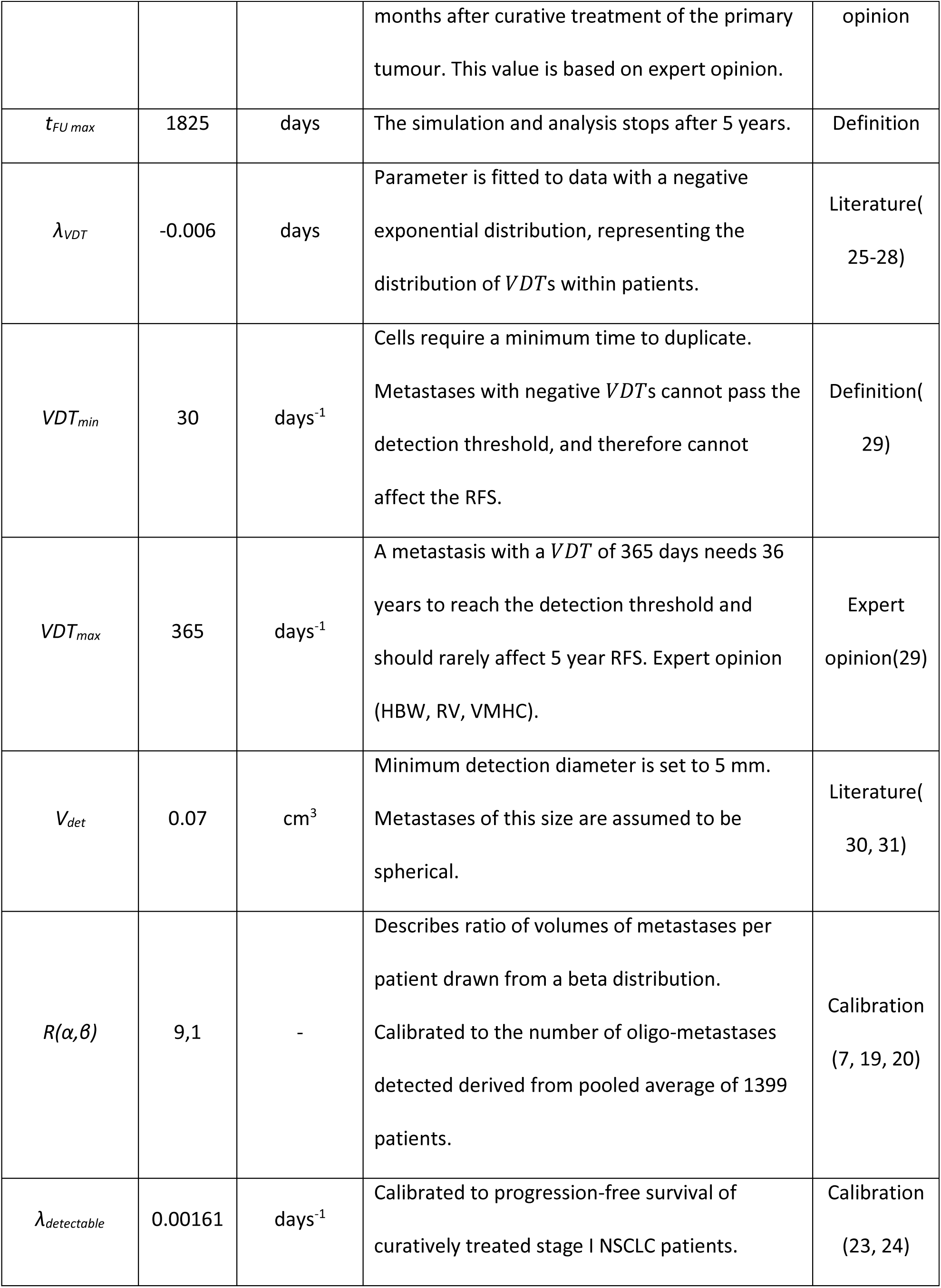
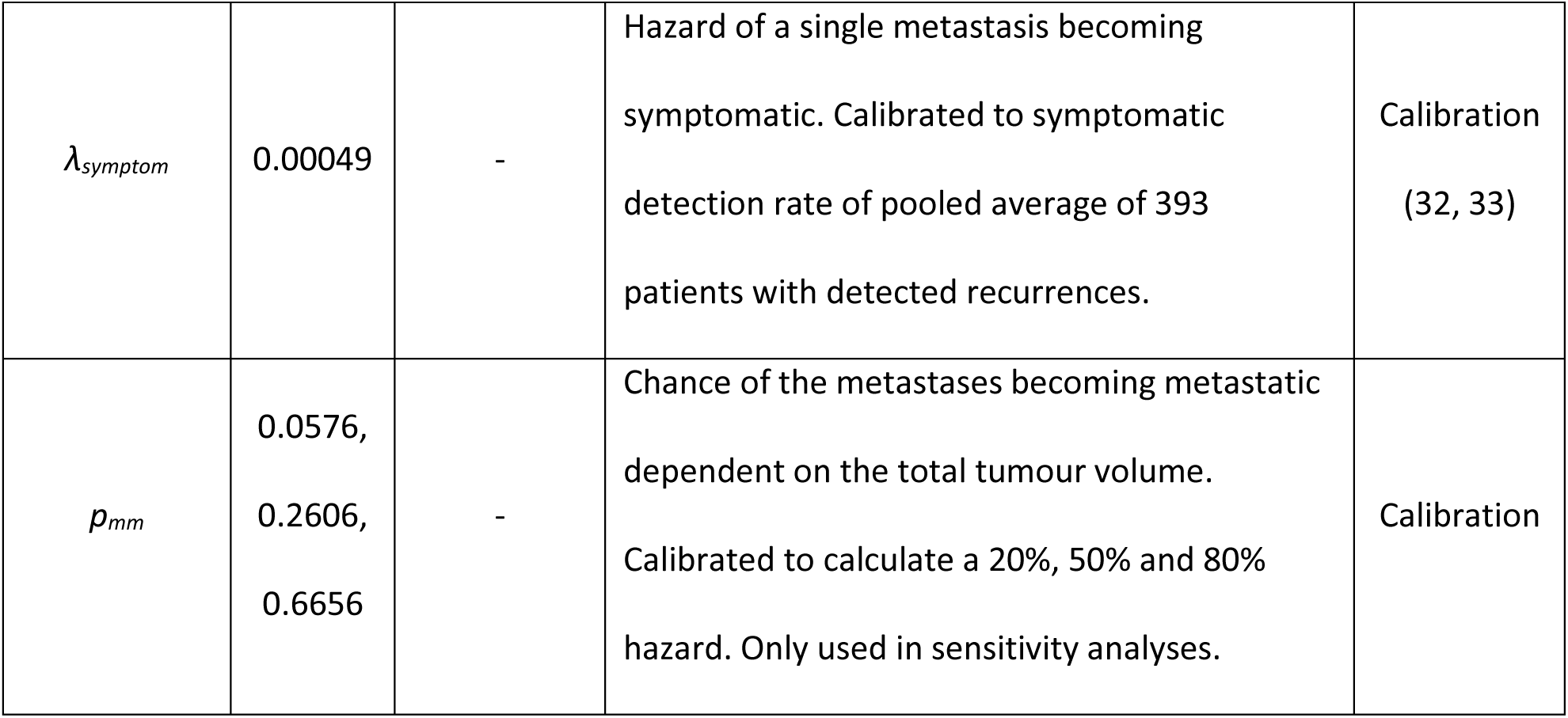
Model Input Parameters

#### Model assumptions

The most important assumptions of the microsimulation-model are listed below, as well as the reasoning behind these assumptions.

1. An exponential tumour growth model using volume doubling time was chosen, because volume doubling time is the most commonly used statistic in literature. All metastases within one patient are assumed to have the same growth rate, and are formed consecutively with fixed time intervals.
2. Each patient is assumed to have a fixed number of metastases after their primary tumour had been curatively treated. The proportion of patients that have zero metastases after curative treatment equals the proportion of patients recurrence-free after 5 years. In all other patients, the number of metastases is randomly drawn from a rounded truncated normal distribution.
3. We assume that metastases below the minimum detectable size for the Computed Tomography scan will always be missed.
4. Metastases of detectable size are either found during surveillance or on an unscheduled scan because of symptoms, whichever happens first. Other scenarios are not considered. Surveillance CT-scans are able to detect recurrences to the lung, liver and adrenal glands. Bone and brain metastases are highly symptomatic. Less than 3% of NSCLC metastases are found in other organs, and these metastases are often also symptomatic(*21, 22*). Therefore, we assume that the combination of symptomatic and surveillance detection sufficiently describe the detection patterns.
5. Once one recurrence is detected, a more rigorous examination, that is, a Positron Emission Tomography–Computed Tomography follows, resulting in detection of all metastases above the minimum detectable size.
6. The proportion of patients with microscopic metastases within those with detected oligo-recurrent disease is assumed to be equal to the proportion of patients with a 5 year PFS after treatment of oligo-recurrent disease.

### Model functions

Each simulated patient starts with a number of metastases *M_total_* drawn from a truncated normal distribution *N*(μ, σ) ≥ 1, rounded up to integer numbers. All metastases in the model grow with a patient specific volume doubling time (*VDT*) in days^−1^.(34)

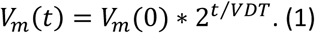

Here *V_m_(t)* is the volume (in cm^3^) of metastasis *m*, *m = 1… M_total_*, at time *t* (in days). The size of the largest metastasis is used to calculate the sizes of the other metastases. Relative sizes of metastases are defined by size ratio *R*:(35)

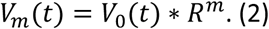

In the model, metastases can either be detected during a routine surveillance scan, or when metastases become symptomatic. Metastases become detectable on a CT-scan once their size is equal or larger than the minimum detectable volume *V_det_*. An exponential hazard function for becoming symptomatic is assumed:

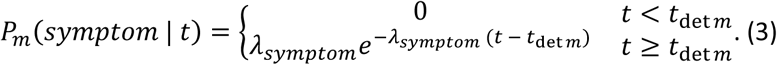

with *λ_symptom_* the hazard to develop symptoms. Time *t_det_* _m_ is defined such that: *V_m_*(*t_det m_*) = *V*_det_. Metastases smaller than *V_det_* are assumed to be too small to cause symptoms.

### Parameter estimation

Model parameters have been estimated from patient data and literature when possible. All unobservable parameters have been estimated using calibration in combination with other observable target parameters.

Data of stage I NSCLC patients curatively treated with video assisted thoracoscopic surgery and stereotactic body radiation therapy was obtained from two studies performed in the Netherlands between 2003 and 2013.(23, 24) Patients were excluded if they had stage ≥ II disease, and ECOG performance score ≥ 2, a second primary tumour or history with previous cancer. Patients of both studies were pooled and 1:1 propensity score matched on treatment using the Matching R package, version 4.9-2. Recurrence Free Survival (RFS) was analysed with a Kaplan Meier survival curve using Statistical Package for Social Sciences version 22.0.

*VDT*s were obtained from papers that reported the distribution of growth rates measured with CT-equipment, in NSCLC with mixed histologies and non-specific tumour morphology.(25–28) Some papers classified growth rates into groups (fast, moderate, slow, no growth). For these groups, the average of the minimum and maximum growth rates was assumed to be the average *VDT* of the reported group. A cumulative exponential distribution was fitted to the reported data using a least squared estimate function (Fig 2). This distribution was used to assign random a *VDT* to patients in the microsimulation-model.

**Fig 2.**
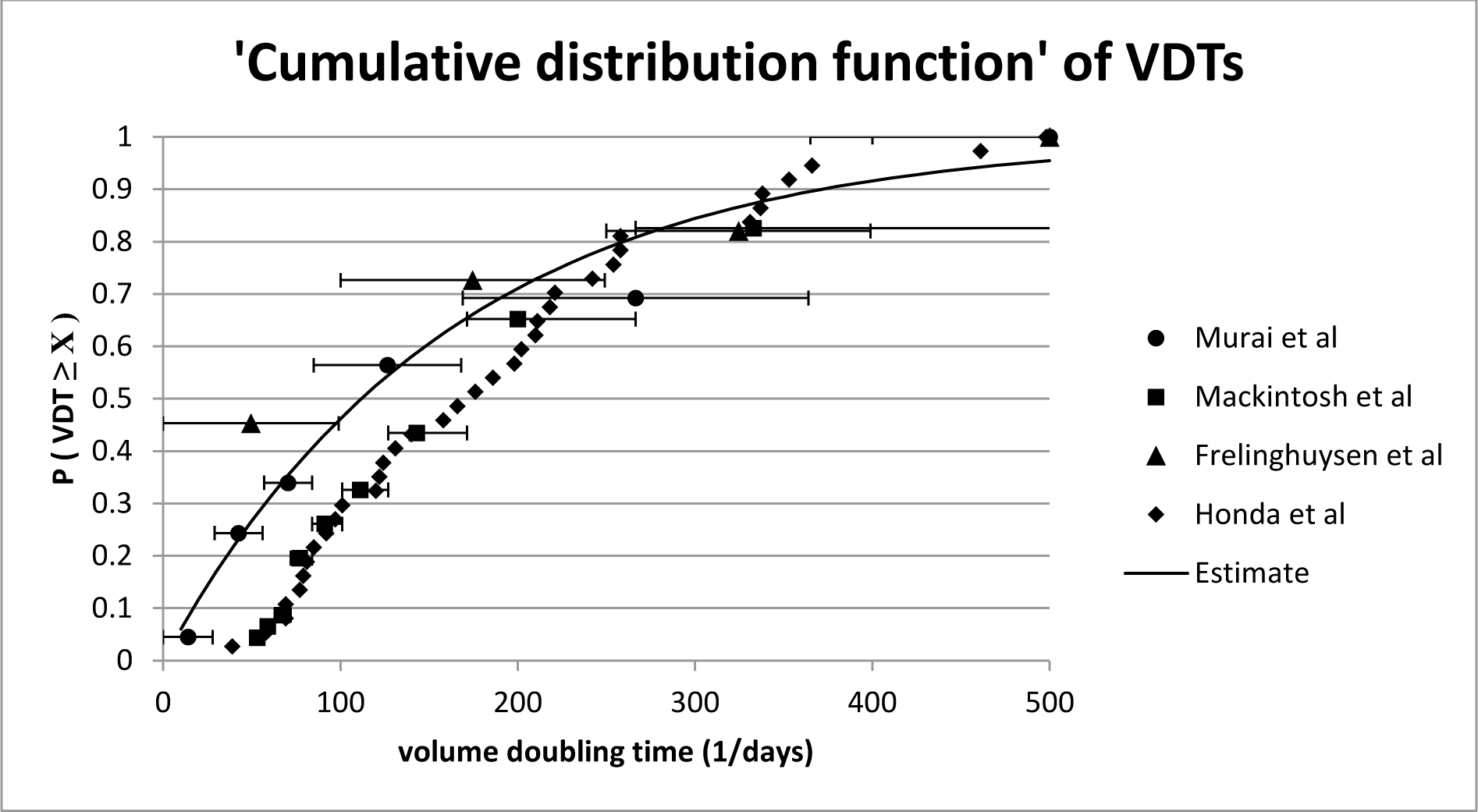
Estimation of a cumulative distribution function of *VDT*. *VDT* values found in literature are used to create a scatterplot.(25–28) A cumulative distribution function is fitted using a least square function to the data shown in this scatterplot to obtain a quantile function, which is subsequently used to determine random tumour growth rates per patient.

Smaller metastases are more likely to be missed on a scan.(36, 37) As such, the reported scan sensitivities and specificities for tumours of any size as found in literature are not suitable for our continuous tumour growth microsimulation-model.(38) Instead, we have therefore chosen to use a detection threshold. We assumed a detection threshold of 5 mm diameter on CT, (30, 31) corresponding with a spherical volume of 0.07 cm^3^.

### Model calibration

Four (sets of) parameters needed to be calibrated: 1) the detection rate parameter *λ_det_*, which is used to randomly draw the time when the largest metastasis passes the detection threshold, 2) the parameters used to determine the total number of metastases per patient *M_total_ (µ,σ)*, 3) the parameters used to determine the size ratio of metastases *R (α,β)*, and 4) the hazard of a metastasis becoming symptomatic (*λ_symptom_*). Each of these model parameters is strongly associated with a specific model output parameter that is also available from literature or patient level data (see below). The mean squared error of 1000 simulations was used in combination with a univariate or grid search algorithm (for *M_total_(µ,σ)*, and *R (α,β)*), and repeated until the calibration target was matched up to 3 significant figure places.(39, 40)

Calibrated parameters only interact in one direction: the number of metastases influences the chance that a patient becomes symptomatic, but the hazard of a single metastasis becoming symptomatic does not influence the number of metastases per patient. Therefore, parameters could be calibrated consecutively if done in the correct order: the most dominant parameter first.

Firstly, the distribution of the time to detectability (that is, reaching a 5mm diameter) of the largest metastasis was assumed to be a negative exponential distribution 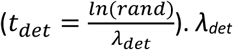 was calibrated against the ½ yearly RFS, weighted for the number of patients at risk. Both the simulations and the patient level data used the same surveillance schedule of the Dutch guidelines.(41)

Secondly, *M_total_* represents the number of both detected and undetectable metastases per patient. There is no data available on this number, or the distribution of detected metastases. Therefore, *M_total_* is assumed to be normally distributed, but is truncated such that *M_total_* ≥ 1. *M_total_ (µ,σ)* was calibrated against the proportion of oligo-metastases without undetected metastases. To determine this proportion, it was assumed that patients who are curatively treated for oligo-metastases and are progression-free at 5 years do not have undetected metastases. A weighted average of estimates reported in the medical scientific was calculated, resulting in an estimated 16% of oligo-metastases that have no underlying undetected metastases.(6–8, 19, 20) To allow a unique solution for *µ* and *σ*, we added to following constraint: the solution for *µ* and *σ* that minimizes the maximum number of detected metastases under the current surveillance schedule.

Thirdly, the size ratio of metastases, *R* is calculated as the volume of the second largest metastasis divided by the volume of the largest metastasis. Therefore, *R* has a value between 0 and 1 by definition. We used a Beta distribution to draw *R* for each patient. The shape parameters *α,β* were defined as integer values. *R(α,β)* was calibrated against the pooled average proportion of oligo-.(7, 19, 20) As a constraint, the solution with the smallest lower tail was selected.

The last calibrated variable was *λsymptom*, the hazard of a single metastasis becoming symptomatic. *λsymptom* was calibrated against the ratio of symptomatically (or unscheduled) detected metastases to metastases detected by a scheduled scan.(32, 33)

### Model simulations

The life histories of 100.000 patients with stage I NSCLC and one or more undetected metastases were simulated from treatment of the primary tumour until the detection of a metastasis (RFS). Patients without micro-metastatic disease after curative treatment of the primary tumour are not simulated, but are added to the cohort post-simulation in accordance to the proportion of patients recurrence-free after 5 years.

For the simulation, each individual was randomly generated from the fitted distributions: the total number of metastases *Mtotal*, the size ratio *R* of metastases, the volume doubling time, the time that the metastases would grow past the detection threshold *tdet* and the time that the patient would develop symptoms *tsymp*. Subsequently, the model determines if symptomatic detection occurs before or after the time that the tumour would be detected on a surveillance scan. Once the time of detection is determined, the number of metastases visible on a scan *mdet* can be calculated with function 4. Function 4 can be derived from function 1 and 2 (Appendix S1). The point in time that the largest metastasis becomes detectable on a scan is denoted by *tdet 0*.

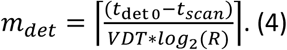

Output of the model contains all the randomly generated patient specific parameters, the size of the largest metastasis and the time and type of detection.

### Prognostic groups

A binary logistic regression model was used for the purpose of identifying clinically observable predictors that can distinguish oligo- from oligo+. The parameters that were considered clinically observable were: the number of recurrences detected, the size category of the largest detected recurrence: small (<6mm), medium (6-8mm), or large (>8mm)(31), symptomatic or surveillance detection, and the RFI(years).(11) Predictors with Wald statistic p < 0.05 were used to divide simulated patients into prognostic sub-groups and determine the proportion of oligo+ for each of these groups. These sub-groups were pooled into a low-risk group with less than 30% oligo+, and a high-risk group with more than 30% oligo+. The 30% risk threshold was considered an acceptable level to consider curative therapy, based on a discussion with our clinical experts (LA, EAK, SYS, and FMNHS). Still, this 30% threshold is suggested tentatively, as it is not obtained from any formal consensus procedure. However, the threshold allows for accessible presentation of the performance and potential use of our modelling approach, and can be seen as a first step towards decision support in this area.

### Sensitivity analyses

To test the robustness of our results, we have designed a series of scenarios and tested how much the predictors used to determine risk groups, the proportion of oligo+ and size of the risk-groups are affected by these scenarios.

1. Recalibration of model parameters to upper and lower confidence interval of their targets.
2. Random variation in *VDT* of metastases within one patient and in the detection threshold.
3. Correlation between the volume doubling time and the total number of metastases per patient.
4. Redefinition of oligo-metastases to 1 or 1 to 5 metastases.
5. The ability of metastases to produce new metastases.

All sensitivity analysis scenarios are described in detail in the Technical Appendix.

## Results

### Calibration

An overview of all calibrated parameters can be found in Table 1.

Fig 3 shows the resulting model-based progression-free survival curve compared to the two Dutch cohorts.(23, 24) The simulated progression-free survival is within the 95% CI range of the cohort data with the exception of the first and third month, but the 95% CI was never exceeded with more than 1.1% after calibration of *λdetectable*.

**Fig 3.**
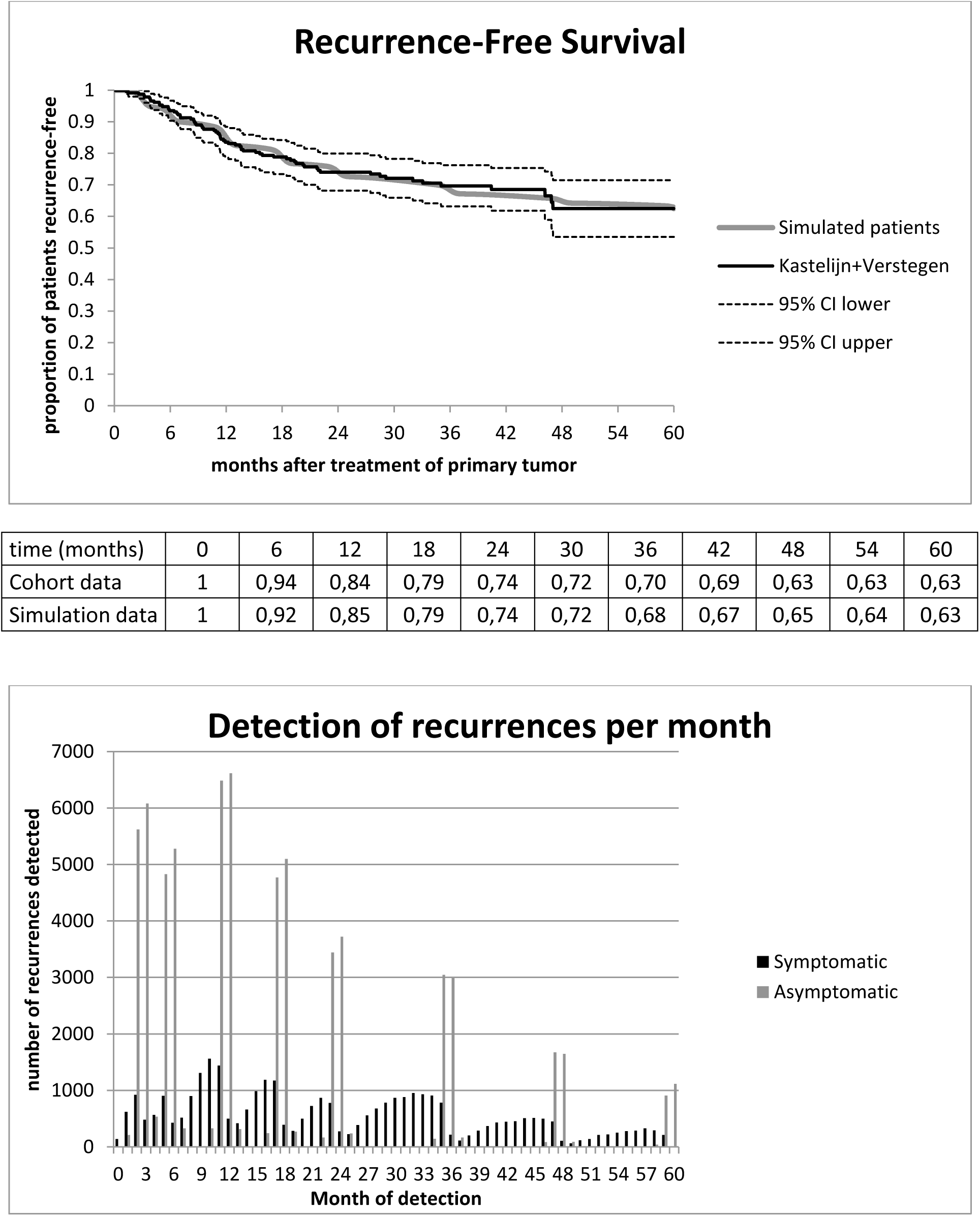
Detection of recurrences. (a) Recurrence-free survival after curative treatment of the primary tumour of simulated patients is calibrated to match two Dutch cohorts.(23, 24) Recurrences are detected with a surveillance scan schedule and unplanned additional scans using a symptom hazard model. 62.5% of the Dutch cohorts remained recurrence-free. Patients without recurrences were not simulated, but have been added after the simulation for the purpose of showing differences between the simulated patients and the Dutch cohorts in this figure. (b) Number of detected recurrences per month. Asymptomatic recurrences are only detected during surveillance scans. These scans are planned according to a fixed schedule, with a small variation in scan date *N*(*μ* = FU *schedule, σ* = 15.5). Symptomatic recurrences are detected throughout the year, and detection times are determined with the hazard model (function 3).

Calibration of *Mtotal* resulted in a maximum of 70, an average of 9, and a median of 6 metastases detected. This solution was considered realistic, based on discussion with our clinical experts LA, EAK, SYS, and FMNHS.

Calibration of size ratio *R* resulted in α = 9 and β = 1, giving an average *R* of 0.93.

The proportion of symptomatically detected metastases was 33.5% after calibration of the parameter *λsymptom*.

### Simulation results

The microsimulation-model was used to produce a patient-level database of 100.000 patients. The first row of Table 2 shows the characteristics for all simulated patients. These patients are subdivided into 3 subgroups in row two to four: poly-metastases (patients with 4 or more detected metastases), oligo+ (patients with 3 or less detected metastases and additional undetectable metastases), and oligo− (patients with 3 or less detected metastases and no additional undetectable metastases).

**Table 2.** Descriptives of generated patient level data (median and 2.5% and 97.5% percentiles)

| | $R$ | $VDT (days^{-1})$ | Diameter<br>(cm) | Metastases<br>detected | Total<br>metastases | RFI (days) |
| --- | --- | --- | --- | --- | --- | --- |
| All patients | 0.93<br>(0.67-1.00) | 103<br>(33-318) | 0.58<br>(0.50-1.13) | 6<br>(1-32) | 21<br>(2-49) | 510<br>(74-1772) |
| Poly<br>metastases | 0.95<br>(0.75-1.00) | 88<br>(33-297) | 0.61<br>(0.51-1.21) | 10<br>(4-35) | 22<br>(5-49) | 542<br>(75-1777) |
| Oligo+ | 0.85<br>(0.59-0.98) | 146<br>(38-337) | 0.53<br>(0.50-0.66) | 2<br>(1-3) | 22<br>(5-49) | 362<br>(71-1579) |
| Oligo- | 0.92<br>(0.67-1.00) | 104<br>(34-319) | 0.59<br>(0.50-1.59) | 2<br>(1-3) | 2<br>(1-3) | 530<br>(77-1817) |
| High-risk<br>group | 0.86<br>(0.60-0.99) | 143<br>(38-336) | 0.53<br>(0.50-0.69) | 2<br>(1-3) | 20<br>(1-49) | 363<br>(72-1629) |
| Low-risk<br>group | 0.91<br>(0.47-1.00) | 55<br>(32-138) | 0.99<br>(0.81-2.95) | 2<br>(1-3) | 2<br>(1-25) | 1075<br>(192-1825) |

Median (2.5% - 97.5% percentile). *VDT*, Volume doubling time; RFI, Recurrence-free interval;
Symptomatic, symptomatic (unscheduled) detection.

Between these three subgroups, the ranges for all reported characteristics are largely overlapping, although the skewness of the distributions is different. The median oligo+ patient has a large size ratio between metastases, slow growing metastases, small diameters at time of detection, and a low RFI.

### Prognostic groups

Although there is a large difference in the median RFI of oligo+ and oligo-groups as shown in Table 2, RFI is not a significant predictor in the logistic regression model (Table 3), when size is included in the model. All other predictors (size, number of detected metastases and symptomatic detection) are significant, and are therefore used to construct Table 4.

**Table 3.** Hazard Ratios predicted by logistic regression model.

| Predictor | Hazard Ratio |
| --- | --- |
| 1 Metastasis detected | reference |
| 2 Metastases detected | 1.76 |
| 3 Metastases detected | 2.44 |
| Asymptomatic detection | reference |
| Symptomatic detection | 1.63 |
| Small size (<6mm) | reference |
| Medium size (6-8mm) | 6.90 |
| Large size (>8mm) | 146.79 |

**Table 4.** Proportion of oligo+ within patients with detected oligo-metastases in the Base Case scenario.

|  | Asymptomatic |  |  | Symptomatic |  |  |
| --- | --- | --- | --- | --- | --- | --- |
| Metastases detected: | 1 | 2 | 3 | 1 | 2 | 3 |
| Small (<6mm) | 0.91 | 0.92 | 0.93 | 0.88 | 0.88 | 0.83 |
| Medium (6-8mm) | 0.08 | 0.61 | 0.81 | 0.08 | 0.71 | 0.84 |
| Large (>8mm) | 0.00 | 0.02 | 0.19 | 0.00 | 0.00 | 0.24 |

Proportions of oligo+ per risk sub-group. Grey fields represents the low-risk group.

A logistic regression model was used as an explorative search for parameters suitable to determine risk groups. Hazard Ratios are calculated as exp(β). Year of detection was removed from the regression model because it was not a significant predictor (Wald test p > 0.05). All other covariates were significant (Wald test p < 0.0001). Note that the significance is influenced by the large number of simulated patients.

Table 4 shows the proportion of oligo+ at time of detection of recurrence within each of the subgroups defined by the three identified predictors. The 30% oligo+ threshold resulted in two distinct groups of patients. Patients with a large sized metastasis or a single medium sized metastasis fall in the low-risk group, and patients with small or 2 or 3 medium sized metastases fall in the high-risk group. Although symptoms do have an effect on the proportion of oligo+, this effect is not large enough to differentiate between low- and high-risk.

The pooled low-risk group has a proportion of 8.1% oligo+, and the high-risk group has a proportion of 89.3% oligo+. As shown in Table 2, the differences between the high- and the low-risk groups are larger than the differences between oligo+ and oligo-. This is a result of the selection procedure for risk groups. The high- and low-risk groups were compared to the ‘treat-all’ and ‘treat-none’ strategies on their strategy performance (Table 5). 86% of patients with an oligo-recurrence are in the oligo+ group which is the cause of the difference in accuracy between ‘treat-all’ and ‘treat-none’.

**Table 5.** Performance of risk-group based treatment selection compared to ‘treat-all’ and ‘treat-none’ strategies.

| Strategy | Low-risk |  | High-risk |  | Performance of chosen strategy (%) |  |  |  |  |
| --- | --- | --- | --- | --- | --- | --- | --- | --- | --- |
|  | Oligo- | Oligo+ | Oligo- | Oligo+ | Sensitivity | Specificity | PPV | NPV | Accuracy |
| Risk groups | 1252 | 111 | 3427 | 28716 | 26.8 | 99.6 | 91.9 | 89.3 | 89.4 |
| Treat-all | 4691 | 28815 | 0 | 0 | 100.0 | 0.0 | 14.0 | 0.0 | 14.0 |
| Treat-none | 0 | 0 | 4691 | 28815 | 0.0 | 100.0 | 0.0 | 86.0 | 86.0 |

For calculation of strategy performance: low-risk oligo- is true positive, low-risk oligo+ is false positive, high-risk oligo- is false negative, and high-risk oligo+ is true negative. PPV, positive predictive value; NPV, negative predictive value.

Additional analysis of the simulation output by extrapolation of the growth model of metastases showed that 73.8% of the oligo+ patients would switch to detectable poly-recurrent disease within 3 months, and 98.2% would become detectable within one year (Appendix Fig S1).

### Sensitivity analyses

A summary of all the sensitivity analyses is given in Table 6. Details of the proportion of oligo+ per subgroup are shown in Appendix Tables S1-S17. The model output such as diameters of metastases, or RFI remained within realistic ranges under all scenarios.

**Table 6.** Proportion of detected oligo-metastases that fall into the risk groups and proportion of oligo+ within the risk groups per scenario.

| Sensitivity analysis scenario | All oligo-metastases |  | Low-risk |  | High-risk |  |
| --- | --- | --- | --- | --- | --- | --- |
|  | N | % Oligo+ | % of all | % Oligo+ | % of all | % Oligo+ |
| Base Case | 33506 | 86.0 | 4.1 | 8.1 | 95.9 | 89.3 |
| $M_{total}$ Lower | 32194 | 91.0 | 2.7 | 11.2 | 97.3 | 93.5 |
| $M_{total}$ Upper | 34506 | 82.0 | 5.1 | 6.2 | 94.9 | 86.3 |
| RFS Lower | 32600 | 85.0 | 4.5 | 7.6 | 95.5 | 88.9 |
| RFS Upper | 34597 | 86.0 | 3.8 | 8.4 | 96.2 | 89.5 |
| $R$ Lower | 28239 | 83.0 | 4.6 | 1.1 | 95.4 | 87.3 |
| $R$ Upper | 35992 | 87.0 | 3.9 | 14.5 | 96.1 | 90.2 |
| Symptomatic Detection Lower | 32917 | 86.0 | 4.2 | 6.3 | 95.8 | 89.2 |
| Symptomatic Detection Upper | 34390 | 86.0 | 4.0 | 7.4 | 96.0 | 89.7 |
| Random Detection Size Scan | 33641 | 86.0 | 3.9 | 9.1 | 96.1 | 89.2 |
| Random $VDT$ | 33721 | 86.0 | 4.2 | 8.9 | 95.8 | 89.0 |
| Correlation $\rho = 0.5$ | 27879 | 83.0 | 7.4 | 6.8 | 92.6 | 89.7 |
| Correlation $\rho = 1.0$ | 23727 | 80.0 | 13.2 | 4.6 | 86.8 | 91.8 |
| Definition of oligo-recurrence = 1* | 12757 | 89.0 | 5.6 | 5.7 | 94.4 | 93.8 |
| Definition of oligo-recurrence = 5* | 47802 | 82.0 | 2.9 | 7.9 | 97.1 | 84.5 |
| Metastatic metastases 20% * | 33506 | 87.2 | 2.1 | 15.1 | 97.9 | 88.8 |
| Metastatic metastases 50% * | 33506 | 89.2 | 1.4 | 21.1 | 98.6 | 90.2 |
| Metastatic metastases 80% * | 33506 | 92.4 | 0.0 | - | 100 | 92.4 |

\*= Risk groups are made up of different subgroups compared to the Base Case.

The classification into two risk groups and the proportion of oligo+ in the low versus high-risk group were similar to our initial analysis in those scenarios in which model parameters were recalibrated (scenarios 1-8). Change in proportion of oligo+ in all simulated patients was associated with change in proportion of oligo+ in the risk groups. This was the most extreme for scenarios in which *Mtotal* and *R* were varied. (Appendix Figs S2A and B).

Random variation in detection size and growth rates of metastases within one patient only caused an small increase of oligo+ patients in the low-risk group (scenarios 9 and 10).

In scenarios 11 and 12, in which model parameters were correlated, the proportion of oligo+ in the low-risk group decreased even though the size of the low-risk group increased. Simultaneously the proportion of oligo+ in the high-risk group increased. Thus, the ability to separate low- from high-risk patients increased with correlated model parameters.

When oligo metastases are defined as 1 metastasis or as 1 to 5 metastases, this change of definition directly affects the total number of patients considered for curative treatment of their oligo-metastases. Furthermore, the number of detected metastases is a predictor that determines if a patient is high-risk or not (Appendix Tables S13 and S14). Using the definitions of oligo-metastasis as a single detected metastasis results in a risk-model with overall better accuracy, but also a smaller number of patients that is considered to have an oligo-recurrence.

Metastatic metastases had a large direct effect on the proportion of oligo+, resulting in a significant increase of the proportion of oligo+ in all subgroups. As such, more subgroups passed the 30% oligo+ threshold, resulting in fewer subgroups to be pooled into the low-risk group. There are no patients with an acceptable risk to undergo curative therapy left, if 80% of all undetectable metastases are metastasizing at the time of curative treatment.

Over the different sensitivity analyses, the proportion of oligo+ at time of detection of an oligo-metastasis remained relatively stable and resulted in acceptable average risks for curative therapy in the low-risk group. Furthermore, the combinations of predictors that were pooled into the low-risk group remained stable with the exception of the metastatic metastases scenarios.

## Discussion

### Key findings

A microsimulation-model of growth and detection of metastases in individual patients was build based on what we considered to be realistic assumptions and an appropriate level of complexity. The model is able to reproduce detection patterns of metastases as seen in the clinical setting, based on growth rates and other model parameters obtained from literature. The model output was used to identify characteristics that allow prediction of the presence of microscopic disease, that is, the presence of additional undetectable metastases in patients with oligo-metastases. Based on three selected risk predictors, the population of patients with an oligo-recurrence could be grouped into a small low-risk group with an 8.1% risk and a high-risk group with 89.3% risk of microscopic disease. Sensitivity analyses showed that the model is robust to changes in parameters.

### Clinical implications

Currently, many stage I NSCLC patients in whom oligo-metastatic disease is detected are curatively treated for these oligo-metastases, although only 16% of those patients remain progression-free for 5 years.(6–8, 19, 20) For these patients, curative treatment provides no benefit but does provide additional risks and side-effects. Multiple prognostic factors for patients with oligo-metastases have been identified, but most of these predictors such as usage of adjuvant chemotherapy are not suitable to predict the presence of undetected metastases and do not allow for selecting those patients that potentially benefit from curative treatment.(11) The key novelty in this paper is that we study the underlying tumour growth to predict which patients are most likely to have additional undetected metastases, rather than identifying prognostic factors for survival.

Current guidelines suggest that curative treatment for oligo-metastases can be considered.(41) As a result, there is considerable variation in treatment of oligo-metastases in clinical practice. As an alternative to a ‘treat-all’ or a ‘treat-none’ strategy, patients can be classified into high-risk and low-risk groups on the basis of prognostic-factors for additional undetected metastases. Low-risk patients could be curatively treated, while high-risk groups could be treated as patients with poly-recurrences.

Clearly, the presented model requires external validation in a clinical setting. Such validation effort seems worthwhile, as our results show that the use of prognostic-subgroups for clinical decision making could potentially reduce incorrect assignment of curative or systemic treatment for patients without or with undetected metastases, compared to the ‘treat-all’ or ‘treat-none’ strategies (Table 5). The use of risk groups is likely to be both cost saving and beneficial for patients, as it may reduce unnecessary harmful treatments. A full cost-benefit analysis, including life expectancy, quality of life, and costs of the treatments would be needed to confirm this assumption.

There are some patient subgroups in the model that have a relatively high chance to be assigned the wrong treatment. Ninety percent of the simulated patients had a very high or low risk of microscopic disease (above 80% or below 10%). A potential management strategy in in the other ten percent could be to offer these patients more frequent surveillance, since the model predicted that 74% of patients with additional microscopic disease would be diagnosed as poly-recurrence within three months (Appendix Fig S2).

Other factors that should be taken into consideration for treatment decision making are the eligibility criteria of a patient for a certain therapy, and the effects of a therapy on symptoms. Therefore, multidisciplinary teams, and shared decision making are important to provide the best treatment for a specific patient. This simulation model is meant as a tool to support in this decision making process.

The microsimulation-model and prognostic groups are not based on assumptions or data of second primary lung cancers, synchronous oligo-metastases, or other types of cancer than NSCLC.

Predictions of risk of undetectable metastases are therefore not necessarily applicable for these patients, although it is possible to collect data on possible predictors such as sizes of metastases in the clinical setting, and test if it is possible to create risk groups for those patients in a similar fashion. It can be informative and of great value for patients to collect clinical data to see if these conclusions are valid in those scenarios.

### Microsimulation-model

Simulation modelling is a useful tool to synthesize available evidence and extrapolate beyond currently observed data, especially in circumstances when relevant information and processes are either not available from a single source or unobservable. With respect to the growth of micro-metastatic lesions, there is a paucity of information that could support medical decision making. However, evidence from cell cultures,(42) mouse models,(43) and observed growth rates in larger tumors(25) provide indications about tumour growth of micro-metastases in real patients. This information can be combined in a simulation model to allow inference on disease aspects important for clinical or policy decisions.

Although this microsimulation-model is useful for improving insight into the metastatic process, and the selection of patients that may benefit from curative therapy, we need to keep in mind that all models are based on various assumptions. We have made an effort to be fully transparent by making all important model assumptions explicit, and presenting the reasoning behind many of the choices made. Some model assumptions could be tested in the sensitivity analyses. However, validation of the model outcomes in the clinical practice is still needed to determine if the model and its assumptions are correct.

### Prognostic groups

Prognostic factors have been identified and used to pool patients into risk groups. The 30% threshold value for low- and high-risk is arbitrary. Other thresholds may be used, and will influence the proportion of undetected metastases in the high- and low-risk groups. This threshold should therefore be in balance with the expected effects on quality of life and survival. Additional cost-effectiveness research can be used to determine the optimal threshold.

The diameter of metastases was found to be the most important prognostic factor in this model. When building the microsimulation-model, the focus was to make the model realistic with an appropriate level of complexity. The diameter predictor was an emergent property of the simulation model, which makes sense theoretically. A large metastasis would require a very large size ratio for the next metastasis to stay under the detection threshold.

The tumour volume can also have a negative impact on survival. For instance Oh et al. reported a 1.04 death hazard ratio for each cm^3^ rise in volume of a brain metastasis.(44) However, in our model all metastases larger than 0.27 cm3 are considered to be large, but these metastases may still be classified as small in the model of Oh et al. Furthermore, the model outcomes are different; Oh et al predict the chance to die from a specific metastasis, while our model predicts the risk of having additional metastases. Thus, patients with larger brain metastases may have a greater chance to benefit from curative treatment and have a greater hazard to die from this metastasis at the same time.

Sensitivity analyses showed a high level of model robustness. Extreme parameter values resulted in minor effects on the ability to predict underlying metastases, and the outcomes still fall within a clinically observable range. Random variation of parameters had little effect on model outcomes. Correlation between the growth rate and the numbers of metastases per patient improved the ability to distinguish between patients with and without additional undetected metastases. We expect that these features are associated, however evidence for this hypothesis is lacking.

Using a different definition of oligo-metastatic disease, that is, detection of only one recurrence affected proportion of undetected metastases in the low-risk group. Extension of the definition to five recurrences did not increase the size of the low-risk group, but would have a negative effect on the relative number of unnecessary curative treatments if the ‘treat all’ strategy was used. These findings match literature in which the number of oligo-metastases per patient is also a prognostic factor.(11)

When increasing the ability of metastases to metastasize, the size of the low-risk group diminishes accordingly. In the case that 80% of the patients have metastatic metastases, no low-risk patients with an indication for curative therapy remain identifiable. Although this model prediction is to be expected, it does not reflect current clinical practice since there are patients that have long progression-free survival after curative treatment of oligo-metastases. Therefore we argue that metastatic metastases may either be rare,(45) or need more than 5 years to become detectable from the single cell state.

### Recommendations on data collection

Most studies on curative treatment for oligo-metastases in NSCLC are based on retrospective data. These retrospective studies are often based on highly selected patients with favourable inclusion criteria, and use varying definitions of oligo-metastases. These studies are therefore susceptible to selection bias.(16, 46) Two randomized studies have been reported with 29 and 49 enrolled patients.(47, 48) Although results show increased progression-free survival, strong evidence for a benefit of curative therapy for oligo-metastatic NSCLC is still absent.

In the absence of high-quality patient level data, we have used microsimulation to provide guidance and generate new hypotheses. To allow validation and improvement or extension of the model, structural reporting of tumour and patient features in a prospective manner would be highly valuable. To validate the model we present here, reporting of the diameters of detected recurrences, affected organs, numbers of recurrences, and mode of detection (by symptoms or not, and which symptoms) is needed. To allow potential further improvement of the model, factors such as genetic markers, histology of the primary tumour, or other prognostic factors(11) may be investigated for this specific purpose. These factors are not included in the current model.

## Supporting information

Technical Appendix

---

**Appendix Fig S1.**
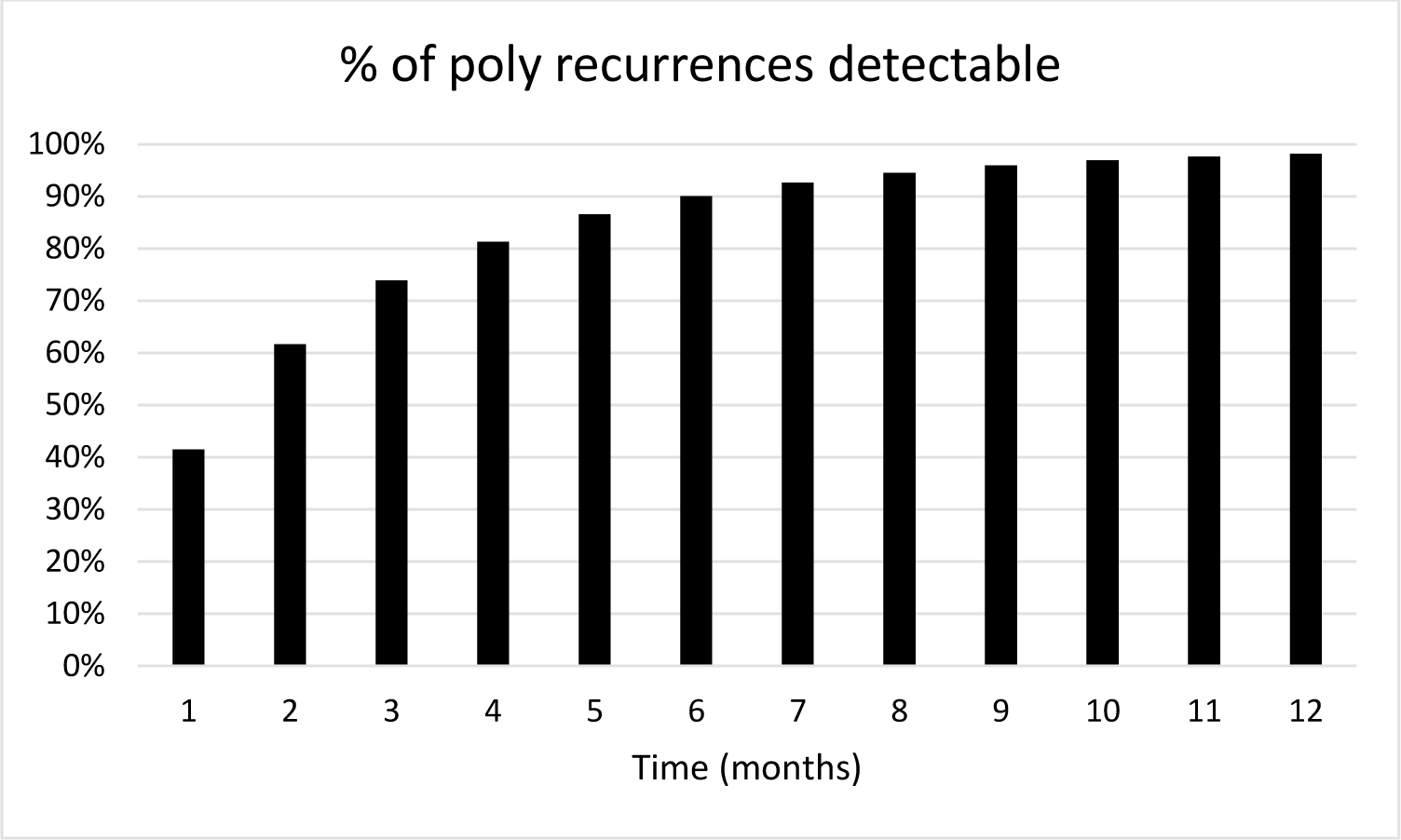
Time required for oligo+ to become distinguishable as poly-recurrent disease. This is estimated as the time that the 4^th^ metastasis passes the detection threshold. 73.8% of oligo+ metastases become detectable after 3 months, and 98.2% of the oligo+ becomes detectable after one year. Data is based on the extrapolation of the microsimulation-model.

**Appendix Fig S2.**
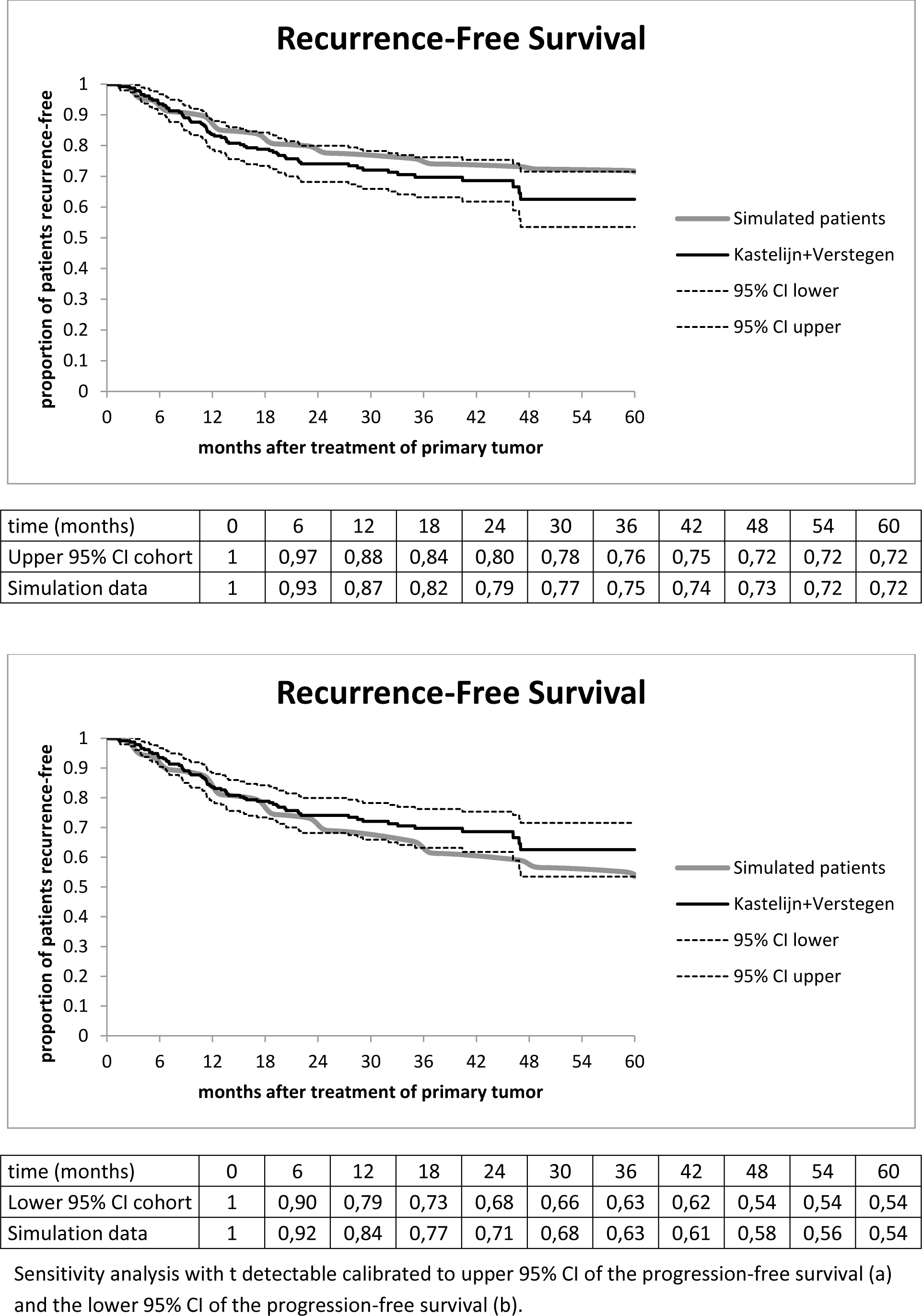
Detection of recurrences in sensitivity analyses. Sensitivity analysis with t detectable calibrated to upper 95% CI of the progression-free survival (a) and the lower 95% CI of the progression-free survival (b).

**Appendix Table S1.** Proportion of oligo+ within patients with detected oligo-metastases in the *Mtotal* Lower scenario.

|  | Asymptomatic |  |  | Symptomatic |  |  |
| --- | --- | --- | --- | --- | --- | --- |
| Metastases detected: | 1 | 2 | 3 | 1 | 2 | 3 |
| Small (<6mm) | 0.95 | 0.95 | 0.96 | 0.91 | 0.91 | 0.89 |
| Medium (6-8mm) | 0.12 | 0.76 | 0.87 | 0.16 | 0.76 | 0.86 |
| Large (>8mm) | 0.00 | 0.02 | 0.25 | 0.00 | 0.06 | 0.27 |

Proportions of oligo+ per risk sub-group. Grey fields represents the low-risk group.

**Appendix Table S2.** Proportion of oligo+ within patients with detected oligo-metastases in the *Mtotal* Upper scenario.

|  | Asymptomatic |  |  | Symptomatic |  |  |
| --- | --- | --- | --- | --- | --- | --- |
| Metastases detected: | 1 | 2 | 3 | 1 | 2 | 3 |
| Small (<6mm) | 0.88 | 0.90 | 0.91 | 0.83 | 0.83 | 0.78 |
| Medium (6-8mm) | 0.06 | 0.58 | 0.77 | 0.07 | 0.58 | 0.73 |
| Large (>8mm) | 0.00 | 0.02 | 0.17 | 0.00 | 0.00 | 0.11 |

Proportions of oligo+ per risk sub-group. Grey fields represents the low-risk group.

**Appendix Table S3.** Proportion of oligo+ within patients with detected oligo-metastases in the RFS Lower scenario.

|  | Asymptomatic |  |  | Symptomatic |  |  |
| --- | --- | --- | --- | --- | --- | --- |
| Metastases detected: | 1 | 2 | 3 | 1 | 2 | 3 |
| Small (<6mm) | 0.91 | 0.92 | 0.91 | 0.89 | 0.86 | 0.83 |
| Medium (6-8mm) | 0.08 | 0.64 | 0.83 | 0.10 | 0.55 | 0.77 |
| Large (>8mm) | 0.00 | 0.02 | 0.17 | 0.00 | 0.06 | 0.19 |

Proportions of oligo+ per risk sub-group. Grey fields represents the low-risk group.

**Appendix Table S4.** Proportion of oligo+ within patients with detected oligo-metastases in the RFS Upper scenario.

|  | Asymptomatic |  |  | Symptomatic |  |  |
| --- | --- | --- | --- | --- | --- | --- |
| Metastases detected: | 1 | 2 | 3 | 1 | 2 | 3 |
| Small (<6mm) | 0.91 | 0.92 | 0.93 | 0.88 | 0.88 | 0.84 |
| Medium (6-8mm) | 0.10 | 0.63 | 0.81 | 0.14 | 0.69 | 0.82 |
| Large (>8mm) | 0.00 | 0.04 | 0.18 | 0.00 | 0.00 | 0.03 |

Proportions of oligo+ per risk sub-group. Grey fields represents the low-risk group.

**Appendix Table S5.** Proportion of oligo+ within patients with detected oligo-metastases in the *R* Lower scenario.

|  | Asymptomatic |  |  | Symptomatic |  |  |
| --- | --- | --- | --- | --- | --- | --- |
| Metastases detected: | 1 | 2 | 3 | 1 | 2 | 3 |
| Small (<6mm) | 0.90 | 0.91 | 0.92 | 0.85 | 0.82 | 0.79 |
| Medium (6-8mm) | 0.01 | 0.40 | 0.69 | 0.00 | 0.41 | 0.69 |
| Large (>8mm) | 0.00 | 0.00 | 0.03 | 0.00 | 0.00 | 0.05 |

Proportions of oligo+ per risk sub-group. Grey fields represents the low-risk group.

**Appendix Table S6.** Proportion of oligo+ within patients with detected oligo-metastases in the *R* Upper scenario.

|  | Asymptomatic |  |  | Symptomatic |  |  |
| --- | --- | --- | --- | --- | --- | --- |
| Metastases detected: | 1 | 2 | 3 | 1 | 2 | 3 |
| Small (<6mm) | 0.92 | 0.93 | 0.93 | 0.87 | 0.86 | 0.85 |
| Medium (6-8mm) | 0.18 | 0.71 | 0.84 | 0.04 | 0.68 | 0.81 |
| Large (>8mm) | 0.00 | 0.05 | 0.28 | 0.00 | 0.07 | 0.22 |

Proportions of oligo+ per risk sub-group. Grey fields represents the low-risk group.

**Appendix Table S7.** Proportion of oligo+ within patients with detected oligo-metastases in the Symptomatic Detection Lower scenario.

|  | Asymptomatic |  |  | Symptomatic |  |  |
| --- | --- | --- | --- | --- | --- | --- |
| Metastases detected: | 1 | 2 | 3 | 1 | 2 | 3 |
| Small (<6mm) | 0.91 | 0.92 | 0.93 | 0.86 | 0.86 | 0.83 |
| Medium (6-8mm) | 0.05 | 0.61 | 0.79 | 0.17 | 0.68 | 0.81 |
| Large (>8mm) | 0.00 | 0.01 | 0.17 | 0.00 | 0.00 | 0.11 |

Proportions of oligo+ per risk sub-group. Grey fields represents the low-risk group.

**Appendix Table S8.** Proportion of oligo+ within patients with detected oligo-metastases in the Symptomatic Detection Upper scenario.

|  | Asymptomatic |  |  | Symptomatic |  |  |
| --- | --- | --- | --- | --- | --- | --- |
| Metastases detected: | 1 | 2 | 3 | 1 | 2 | 3 |
| Small (<6mm) | 0.91 | 0.93 | 0.93 | 0.88 | 0.87 | 0.84 |
| Medium (6-8mm) | 0.07 | 0.65 | 0.82 | 0.13 | 0.68 | 0.81 |
| Large (>8mm) | 0.00 | 0.01 | 0.20 | 0.00 | 0.03 | 0.18 |

Proportions of oligo+ per risk sub-group. Grey fields represents the low-risk group.

**Appendix Table S9.** Proportion of oligo+ within patients with detected oligo-metastases in the Random Detection Size Scan scenario.

|  | Asymptomatic |  |  | Symptomatic |  |  |
| --- | --- | --- | --- | --- | --- | --- |
| Metastases detected: | 1 | 2 | 3 | 1 | 2 | 3 |
| Small (<6mm) | 0.91 | 0.92 | 0.92 | 0.84 | 0.89 | 0.84 |
| Medium (6-8mm) | 0.10 | 0.66 | 0.80 | 0.07 | 0.71 | 0.80 |
| Large (>8mm) | 0.00 | 0.02 | 0.22 | 0.00 | 0.03 | 0.25 |

Proportions of oligo+ per risk sub-group. Grey fields represents the low-risk group.

**Appendix Table S10.** Proportion of oligo+ within patients with detected oligo-metastases in the Random *VDT* scenario.

|  | Asymptomatic |  |  | Symptomatic |  |  |
| --- | --- | --- | --- | --- | --- | --- |
| Metastases detected: | 1 | 2 | 3 | 1 | 2 | 3 |
| Small (<6mm) | 0.90 | 0.92 | 0.93 | 0.89 | 0.86 | 0.83 |
| Medium (6-8mm) | 0.11 | 0.64 | 0.80 | 0.00 | 0.68 | 0.83 |
| Large (>8mm) | 0.00 | 0.02 | 0.19 | 0.00 | 0.05 | 0.18 |

Proportions of oligo+ per risk sub-group. Grey fields represents the low-risk group.

**Appendix Table S11.** Proportion of oligo+ within patients with detected oligo-metastases in the Correlation ρ = 0.5 scenario.

|  | Asymptomatic |  |  | Symptomatic |  |  |
| --- | --- | --- | --- | --- | --- | --- |
| Metastases detected: | 1 | 2 | 3 | 1 | 2 | 3 |
| Small (<6mm) | 0.93 | 0.93 | 0.94 | 0.94 | 0.90 | 0.85 |
| Medium (6-8mm) | 0.10 | 0.65 | 0.80 | 0.15 | 0.57 | 0.81 |
| Large (>8mm) | 0.00 | 0.02 | 0.14 | 0.00 | 0.00 | 0.11 |

Proportions of oligo+ per risk sub-group. Grey fields represents the low-risk group.

**Appendix Table S12.** Proportion of oligo+ within patients with detected oligo-metastases in the Correlation ρ = 1.0 scenario.

|  | Asymptomatic |  |  | Symptomatic |  |  |
| --- | --- | --- | --- | --- | --- | --- |
| Metastases detected: | 1 | 2 | 3 | 1 | 2 | 3 |
| Small (<6mm) | 0.95 | 0.95 | 0.96 | 0.93 | 0.92 | 0.90 |
| Medium (6-8mm) | 0.11 | 0.69 | 0.83 | 0.15 | 0.64 | 0.86 |
| Large (>8mm) | 0.00 | 0.01 | 0.09 | 0.00 | 0.00 | 0.09 |

Proportions of oligo+ per risk sub-group. Grey fields represents the low-risk group.

**Appendix Table S13.**
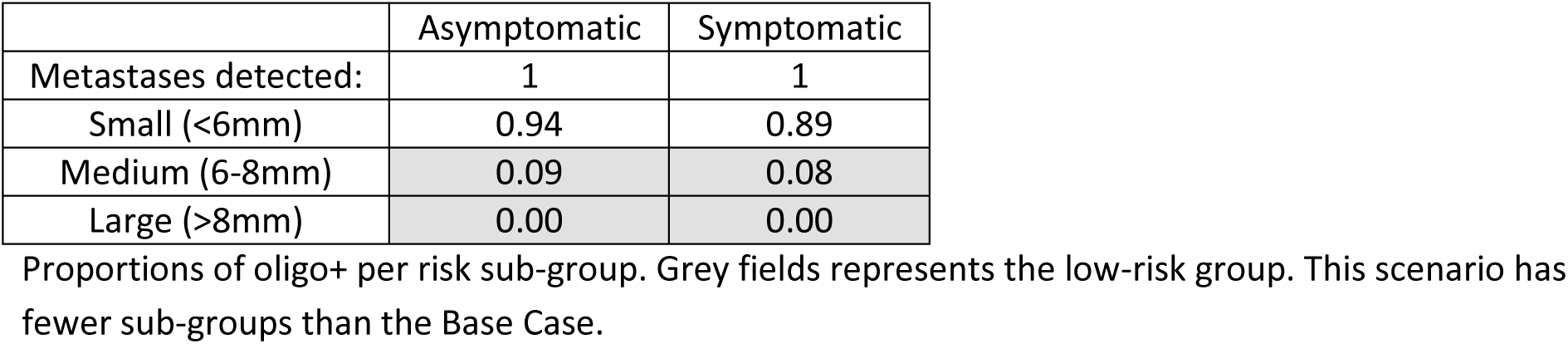
Proportion of oligo+ within patients with detected oligo-metastases in the redefinition of oligo metastases to 1 metastasis scenario.

|  | Asymptomatic | Symptomatic |
| --- | --- | --- |
| Metastases detected: | 1 | 1 |
| Small (<6mm) | 0.94 | 0.89 |
| Medium (6-8mm) | 0.09 | 0.08 |
| Large (>8mm) | 0.00 | 0.00 |

Proportions of oligo+ per risk sub-group. Grey fields represents the low-risk group. This scenario has fewer sub-groups than the Base Case.

**Appendix Table S14.**
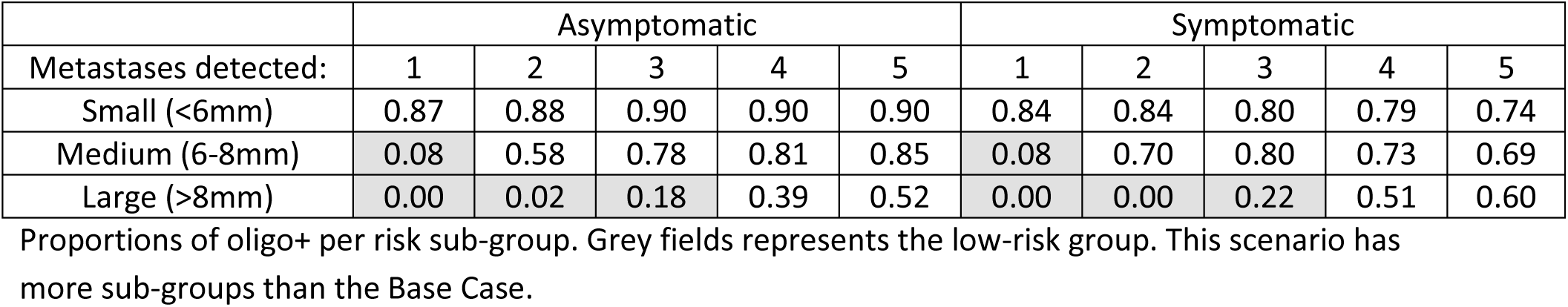
Proportion of oligo+ within patients with detected oligo-metastases in the redefinition of oligo metastases to 1-5 metastases scenario.

|  | Asymptomatic |  |  |  |  | Symptomatic |  |  |  |  |
| --- | --- | --- | --- | --- | --- | --- | --- | --- | --- | --- |
| Metastases detected: | 1 | 2 | 3 | 4 | 5 | 1 | 2 | 3 | 4 | 5 |
| Small (<6mm) | 0.87 | 0.88 | 0.90 | 0.90 | 0.90 | 0.84 | 0.84 | 0.80 | 0.79 | 0.74 |
| Medium (6-8mm) | 0.08 | 0.58 | 0.78 | 0.81 | 0.85 | 0.08 | 0.70 | 0.80 | 0.73 | 0.69 |
| Large (>8mm) | 0.00 | 0.02 | 0.18 | 0.39 | 0.52 | 0.00 | 0.00 | 0.22 | 0.51 | 0.60 |

Proportions of oligo+ per risk sub-group. Grey fields represents the low-risk group. This scenario has more sub-groups than the Base Case.

**Appendix Table S15.** Proportion of oligo+ within patients with detected oligo-metastases in the Metastatic metastases 20% scenario.

|  | Asymptomatic |  |  | Symptomatic |  |  |
| --- | --- | --- | --- | --- | --- | --- |
| Metastases detected: | 1 | 2 | 3 | 1 | 2 | 3 |
| Small (<6mm) | 0.91 | 0.92 | 0.94 | 0.88 | 0.88 | 0.84 |
| Medium (6-8mm) | 0.12 | 0.64 | 0.83 | 0.14 | 0.73 | 0.85 |
| Large (>8mm) | 0.21 | 0.34 | 0.48 | 0.16 | 0.32 | 0.46 |

Proportions of oligo+ per risk sub-group. Grey fields represents the low-risk group. This scenario has fewer categories that are considered low risk than the Base Case.

**Appendix Table S16.**
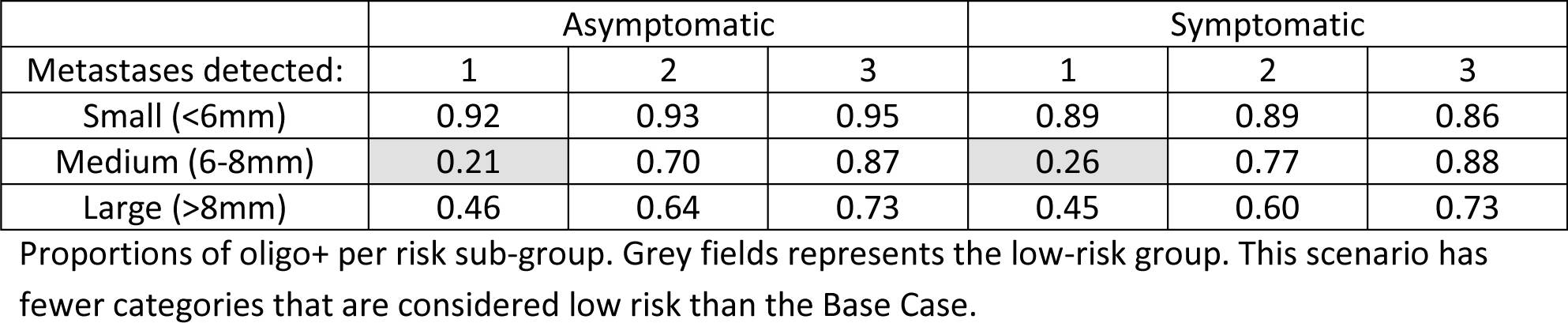
Proportion of oligo+ within patients with detected oligo-metastases in the Metastatic metastases 50% scenario.

**Appendix Table S17.** Proportion of oligo+ within patients with detected oligo-metastases in the Metastatic metastases 80% scenario.

|  | Asymptomatic |  |  | Symptomatic |  |  |
| --- | --- | --- | --- | --- | --- | --- |
| Metastases detected: | 1 | 2 | 3 | 1 | 2 | 3 |
| Small (<6mm) | 0.93 | 0.95 | 0.96 | 0.90 | 0.91 | 0.90 |
| Medium (6-8mm) | 0.40 | 0.81 | 0.93 | 0.45 | 0.86 | 0.93 |
| Large (>8mm) | 0.74 | 0.89 | 0.93 | 0.77 | 0.85 | 0.93 |

Proportions of oligo+ per risk sub-group. This scenario has no low-risk categories in contrast to the Base Case.

