## Supplementary material for "Prediction of microscopic metastases in patients with metachronous oligo-metastases after curative treatment of Non-Small Cell Lung Cancer": Technical Appendix

**S1 Derivation of function 4**

$m_{det}$ is the number of detected metastases, and is per definition an integer number between 0 and $m_{total}$. The number of detected metastases is equal to the number of metastases of detectable size ($V_{det}$) at the time that a scan occurs ($t_{scan}$). Metastasis *i* reaches detectable size ($V_{det}$) at $t_{det\,i}$, thus *i* metastases are detectable when $t_{\det i+1} \geq t_{scan} \geq t_{\det i}$. Furthermore, the detection threshold should also fall in between $V_i$ and $V_{i+1}$ at the time of this scan: $V_{i+1}(t_{scan}) \leq V_{det} \leq V_i(t_{scan})$. $m_{det}$ can be obtained by solving *i*.

Applying function 1 on the formula above results in:

$$V_{i+1}(0) * 2^{t_{scan}/VDT} \leq V_0(0) * 2^{t_{\det 0}/VDT} \leq V_i(0) * 2^{t_{scan}/VDT}.$$

And according to function 2:

$$V_0(0) * R^{i+1} * 2^{t_{scan}/VDT} \leq V_0(0) * 2^{t_{\det 0}/VDT} \leq V_0(0) * R^{i} * 2^{t_{scan}/VDT}.$$

This can be simplified to: $i + 1 \leq \frac{(t_{\det 0} - t_{scan})}{VDT * log_2(R)} \leq i.$

$m_{det}$ can be obtained by rounding $i$ to an integer number: $\left\lceil \frac{(t_{\det 0} - t_{scan})}{VDT * log_2(R)} \right\rceil = m_{det}.$

**S2 Sensitivity analyses**

1. *Alternative calibration targets*

The accuracy of the risk groups was tested by generating a new patient database using the regular prediction model with extreme parameters.

For each of the four model parameters that were obtained by calibration, one by one, the upper and lower limit of the 95% confidence intervals around the calibration targets were used to recalibrate the model. For pooled averages of rates (μ) from literature (for oligo-metastases detected, oligo-, and symptomatic detections) the 95% confidence interval around this rate was estimated to be $\mu \pm 1.96\sqrt{\frac{\mu\,(1-\mu)}{n}}$. These recalibrated microsimulation-model were run again and the risk sub-group analyses were repeated. All the other model parameters were left unchanged.

2. *Random normal variation around model parameters*

In the second set of sensitivity analyses, the parameters $VDT$ and $V_{det}$ were separately varied around the point estimates by randomly drawing a new value from a normal distribution per patient. The scan detection threshold, was altered around 5mm with a standard deviation of 1 mm. The individually drawn volume doubling time was altered for the smallest detectable metastases with a standard deviation of 10 days. The microsimulation-model was run again and the risk sub-group analyses were repeated. All the other model parameters were left unchanged.

*3. Correlation between volume doubling time and the total number of metastases*

In the base-case model, no interactions were assumed between model parameters, because proof for such interactions was lacking. However, it is a credible assumption to make that patients with higher *VDT*s also have higher numbers of metastases. We explored the effect of such an interaction by adding a correlation between $VDT$ and $M_{total}$ using the formula:

$$VDT = \frac{\ln\left(\rho * \frac{(M_{total} - \mu_{M_{total}})}{\sigma_{M_{total}}} + (1-\rho) * r\right)}{\lambda_{VDT}}. \quad (5)$$

With $\rho$ as the 'correlation coefficient', and $r$ as a random uniform value that is drawn to determine the $VDT$ of a patients metastases. Two simulations were used for the sensitivity analyses with $\rho = 0.5$ and $\rho = 1.0$. The microsimulation-model was run again and the risk sub-group analyses were repeated. All the other model parameters were left unchanged.

*4. Adaptation of the definition of oligo-metastases*

For these sensitivity analyses, the simulated patient level database was unaltered, but the prediction model was modified by setting the maximum number of detected recurrences that is considered to be an oligo-metastasis $M_{oligo\ max}$ to 1 and to 5. This change in definition, alters the number of patients with detected oligo-metastases and the proportions of oligo+ and oligo- within the risk sub-groups.

*5. Metastasizing metastases*

A sensitivity analysis was included that estimates the impact of the assumption that metastases are able to produce additional metastases, without the presence of the primary tumour. This increases the proportion of oligo+. For this sensitivity analysis, we assumed that the mutations required for the metastasis to metastasize are related to the number of DNA duplication steps and cell divisions.(46) Therefore, we assume that the hazard of metastases acquiring the ability to metastasize is related to the total tumour volume in time:

$$Hazard = 1 - (1 - p_{mm})^{V_{total}(t)}. \quad (6)$$

Here $p_{mm}$ is the probability that one metastatic cell within the total tumour volume will acquire the ability to disseminate metastases, and $V_{total}(t)$ is the total tumour volume in time. The hazard was used to estimate how many of the patients with oligo-metastases in the model have developed additional metastases after removal of the primary tumour. $p_{mm}$ was calibrated to create scenarios with 20%, 50% and 80% metastatic metastases (Table 1). The proportions of oligo+ per risk sub-group was subsequently adjusted for patients that switched from oligo- to oligo+.
